# Gut microbiota composition explains more variance in the host cardiometabolic risk than genetic ancestry

**DOI:** 10.1101/394726

**Authors:** Sandra J. Guzmán-Castañeda, Esteban L. Ortega-Vega, Jacobo de la Cuesta-Zuluaga, Eliana P. Velásquez-Mejía, Winston Rojas, Gabriel Bedoya, Juan S. Escobar

**Affiliations:** Grupo de Investigación en Genética Molecular (GENMOL), Sede de Investigación Universitaria, Universidad de Antioquia, Medellin, Colombia; Vidarium–Nutrition, Health and Wellness Research Center, Grupo Empresarial Nutresa, Medellin, Colombia; Max Planck Institute for Developmental Biology, Tübingen, Germany

**Keywords:** Ancestry informative markers, genetic admixture, mestizo, Latin America, gut microbiota, OTU, non-communicable diseases

## Abstract

**Background:** Cardiometabolic affections greatly contribute to the global burden of disease. The susceptibility to these conditions associates with the ancestral genetic composition and gut microbiota. However, studies explicitly testing associations between genetic ancestry and gut microbes are rare. We examined whether the ancestral genetic composition was associated with gut microbiota, and split apart the effects of genetic and non-genetic factors on host health.

**Results:** We performed a cross-sectional study of 441 community-dwelling Colombian mestizos from five cities. We characterized the host genetic ancestry using 40 ancestry informative markers and gut microbiota through 16S rRNA gene sequencing. We measured variables related to cardiometabolic health (adiposity, blood chemistry and blood pressure), diet (calories, macronutrients and fiber) and lifestyle (physical activity, smoking and medicament consumption). The ancestral genetic composition of the studied population was 67±6% European, 21±5% Native American and 12±5% African. While we found limited evidence of associations between genetic ancestry and gut microbiota or disease risk, we observed a strong link between gut microbes and cardiometabolic health. Multivariable-adjusted linear models indicated that gut microbiota was more likely to explain variance in host health than genetic ancestry. Further, we identified 9 OTUs associated with increased disease risk and 11 with decreased risk.

**Conclusions:** Gut microbiota seems to be more meaningful to explain cardiometabolic disease risk than genetic ancestry in this mestizo population. Our study suggests that novel ways to control cardiometabolic disease risk, through modulation of the gut microbial community, could be applied regardless of the genetic ancestry of the intervened population.

## Background

Obesity, cardiovascular disease and type 2 diabetes are notable contributors to the global burden of disease [1]. Seminal studies in monozygotic twins demonstrated that these cardiometabolic diseases are heritable [2–4], but genome-wide association studies (GWAS) have failed to consistently uncover replicable variants across human populations, with notable exceptions [5,6]. One possible explanation for this is that the identification of variants in candidate genes is highly dependent on the ethnic and geographic origin of the studied population [7]. Differences in allele frequencies and linkage disequilibrium structure make difficult the extrapolation of results between human groups with different genetic backgrounds. Therefore, the ancestral genetic composition of the studied population becomes a key element in association studies [8].

Additionally, the lack of replicability of many GWAS results across populations may be explained by the interactions between gene variants and non-genetic factors affecting the aforementioned complex phenotypes [9]. The gut microbiota, the set of microorganisms that naturally colonize the human intestine [10], is one of such factors. The gut microbiota has been shown to be central to host health [11–13], and to be shaped by human genetics [14,15]. Despite the impact of recent discoveries on the relationship between gut microbes and human health, the degree to which associations found in one population can extend to another is still unclear. The geographic origin of human populations is one of the most important factors shaping the composition of this microbial community [16,17], yet it is unknown whether such biogeographic pattern is explained by host genetics or by non-genetic factors correlated with geography and ancestry (*e.g.*, diet, lifestyle). Studies explicitly testing associations between host genetic ancestry and gut microbiota are still very rare [18].

In this study, we analyzed a cohort of Colombian adults, whose genetic background is product of extensive recent admixture between three continental populations: Europeans, Native Americans and Africans [19]. In these individuals, we estimated the ancestral genetic composition with ancestry informative markers (AIMs), characterized gut microbiota through high-throughput 16S rRNA gene sequencing and measured numerous variables that inform about diet, lifestyle and cardiometabolic disease risk. We aimed to determine whether the ancestral genetic composition of this population was associated with the structure of the gut microbiota, and split apart the effects of genetic and non-genetic factors on human health.

## Results

### The city of origin accounts for differences in the ancestral genetic composition

We performed a cross-sectional study in which we enrolled 441 adult Colombian mestizos in roughly similar proportions across five cities spanning the Colombian Andes and both its Caribbean and Pacific coasts (Bogota, Medellin, Cali, Barranquilla and Bucaramanga); body mass index (BMI: lean, overweight, obese); sex (male, female); and age range (18-40 years, 41-62 years). We characterized the ancestral genetic composition in 440 of these participants using a panel of 40 ancestry informative markers (AIMs) that have been previously shown to discriminate among European, Native American and African populations [20,21] (Table S1). One individual of our cohort could not be genotyped because we were not able to acquire DNA from blood. Overall, the 40 evaluated AIMs were found in Hardy-Weinberg equilibrium (all p>0.05 in exact Hardy-Weinberg tests).

Overall, the ancestral genetic composition of the individuals of this cohort was (mean ± SD) 0.674 ± 0.057 European (range: 0.469–0.788); 0.209 ± 0.048 Native American (0.089– 0.397); and 0.117 ± 0.047 African (0.051–0.352) (Figure 1A). These values differed significantly among cities (ANOVA for European: F_4,435_ = 2.94, p = 0.02; Native American: F_4,435_ = 7.69, p<0.0001; African: F_4,435_= 5.78, p = 0.0002): the European component was highest in Medellin (Northwestern Andes) and lowest in Barranquilla (Northern Caribbean); the Native American component highest in Bogota (Central Andes) and lowest in Medellin; and the African component highest in Barranquilla and lowest in Bogota (Figure 1B–1D). In agreement with this, we found evidence of limited but significant genetic structure (mean *F*_st_ ± SE = 0.004 ± 0.001, 95% CI = 0.002–0.006). However, there was no evidence of isolation by distance, according to a Mantel test considering genetic (*F*_st_/(1-*F*_st_)) and log-transformed geographic distance matrices (r= −0.43, 95% CI = −0.94–0.14, two-tailed p = 0.43). Furthermore, we did not find significant differences in the ancestral genetic composition by other factors controlled by design (p>0.10 in all ANOVAs for BMI, sex and age range).

**Figure 1.**
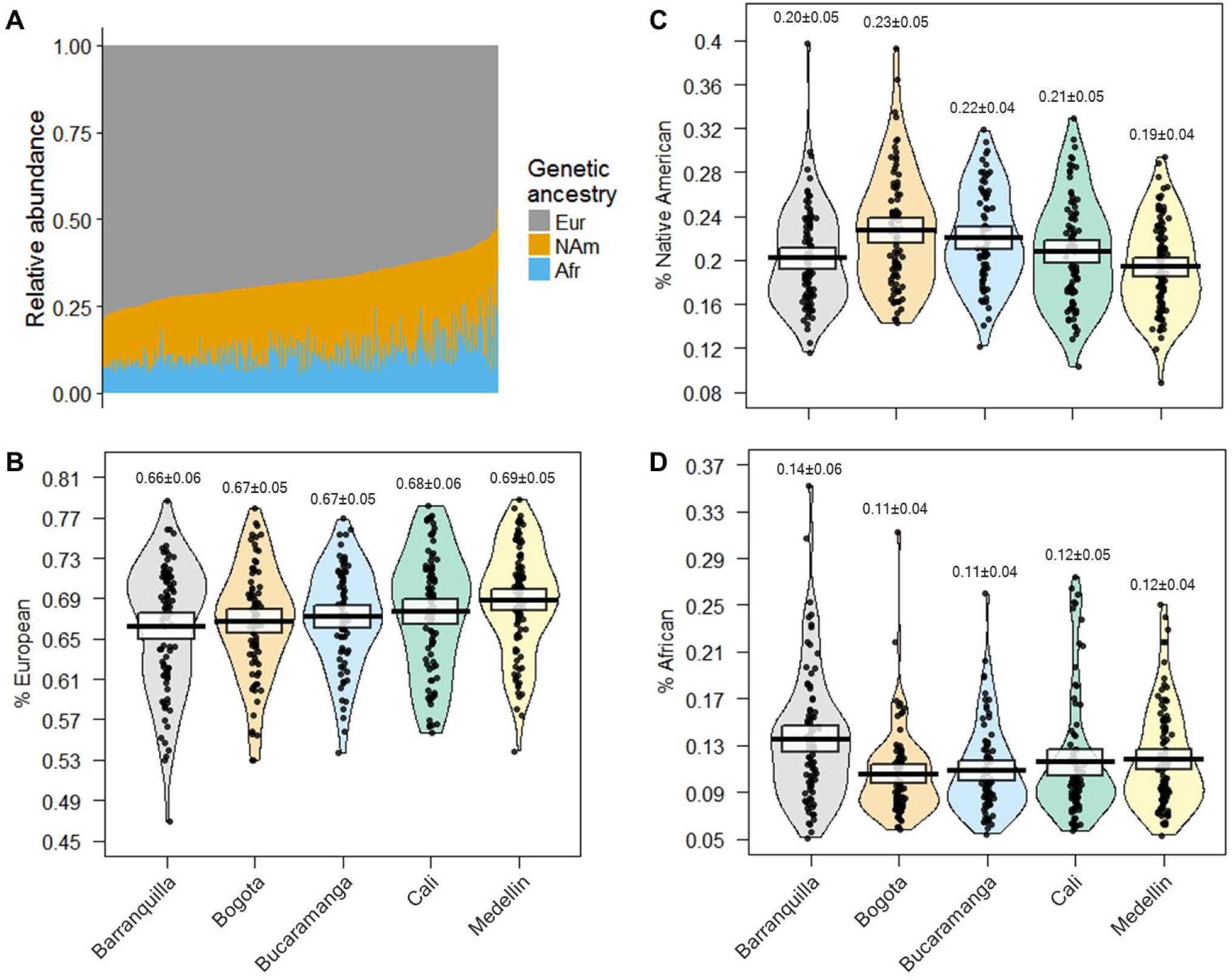
Contributions of European, Native American and African ancestries to the studied population. (A) Ancestral genetic composition across individuals (vertical bars). Data sorted by European component. Eur=European; NAm=Native American; Afr=African. (B-D) Ancestral genetic composition along the five Colombian cities from which participants originated. The raw data, average and 95% confidence intervals are shown in each plot. Mean ± SD given above each plot. Note the change in scale among panels.

Next, we performed a robust principal component analysis (PCA) for compositional data based on the individual proportions of European, Native American and African, and found a gradient where the first component (PC1) distinguished Native American and African ancestries, whereas the second component (PC2) discerned European and non-European ancestries (Figure 2A-2C). In accordance with our previous result, these two components differed among the cities from which participants originated (ANOVA for PC1: F_4,435_ = 7.60, p<0.0001; PC2: F_4,435_ = 3.63, p = 0.006) but did not differ by BMI, sex or age range (p>0.10 in all ANOVAs).

**Figure 2.**
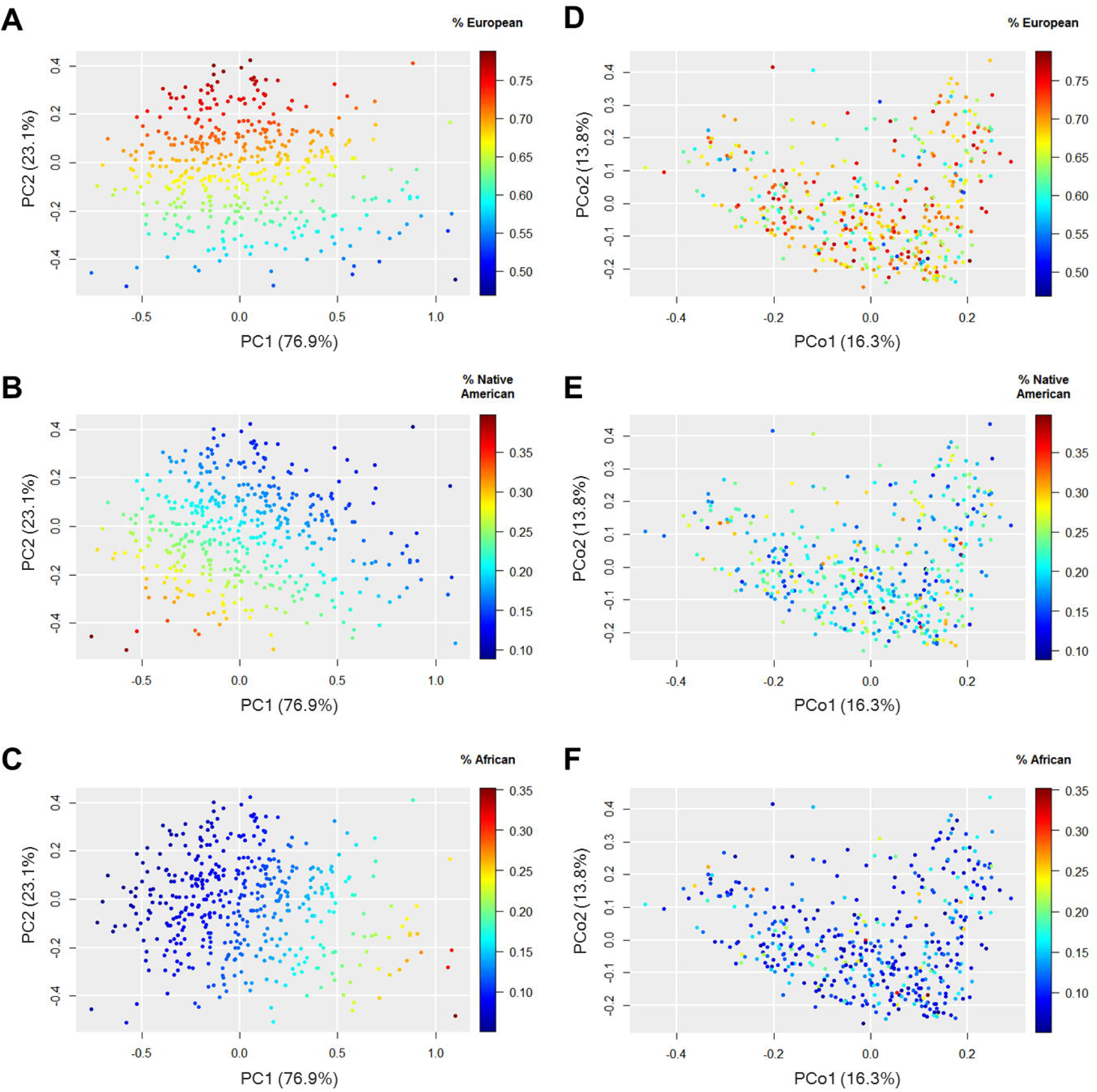
Ancestral genetic composition and gut microbiota composition in the studied population. Each set of panels shows the same cloud point colored by the contributions of each ancestry: (A–C) Robust principal components analysis (PCA) for compositional data based on the proportions of European (A), Native American (B) and African (C) ancestries. (D–F) Principal coordinate analysis (PCoA) based on weighted UniFrac distances of gut microbiota for European (D), Native American (E), and African (F) ancestries. Percentages on the axes represent the proportion of explained variation. Note the change in scale among panels.

### Limited evidence of an association between host genetic ancestry and gut microbiota

Afterwards, we sought to examine whether the host genetic ancestry associated with the composition of gut microbiota. We analyzed the complete microbial community through principal coordinate analysis (PCoA) using weighted UniFrac distances on rarefied sequence counts, and found that the gut microbiota of Colombians formed a single point cloud of microbial abundances. Beta-diversity analyses indicated that differences in the structure of the microbial community were partly driven by the city of origin (PERMANOVA: R^2^ = 0.074, p = 0.001), BMI (R^2^ = 0.010, p = 0.003) and sex (R^2^ = 0.012, p = 0.001), but not by the age range (R^2^ = 0.003, p = 0.22). It is noteworthy that despite the significant association of gut microbiota with the city of origin, we did not find convincing evidence of a direct association between the microbial community and the host genetic ancestry, as shown by Procrustes analyses correlating weighted UniFrac distances and the individual proportions of European, Native American and African (Procrustes correlation = 0.04, p = 0.99), or the first two PCoA axes and PCA components (Procrustes correlation = 0.03, p = 0.91) (Figure 2D-2F).

We further examined whether specific groups of microbes were associated with the host genetic ancestry. First, we correlated the ancestral genetic composition (either as individual proportions of European, Native American and African, or as genetic PCA components) and the relative abundances of dominant taxonomic ranks. At the family level, we observed a positive correlation between the relative abundance of *Enterococcaceae* (*Firmicutes*) and genetic PC1 (Spearman’s rho = 0.16, p = 0.001, q = 0.17), meaning that this family was more abundant in individuals with higher contribution of African ancestry. We did not observe other significant correlations between the host genetic ancestry and relative abundances at the phylum, class, order, genus or species levels (q >0.20; Table S2).

Next, we performed a similar analysis using the relative abundances of operational taxonomic units (OTUs) instead of taxonomic ranks. In this case, we restricted the comparisons to the 100 most abundant OTUs, whose median relative abundances were ≥0.01% across all samples and which comprised up to 80 ± 12% of all 16S rRNA gene reads, thus minimizing potential artifacts produced by sequencing errors. We found that the relative abundance of Otu00068 (*Enterococcus casseliflavus*) directly correlated with genetic PC1 (rho = 0.15, p = 0.001, q = 0.12), corroborating the association described above. No other OTU associated with the host genetic ancestry (q>0.20; Table S3).

Non-genetic factors intimately associated with the host genetic composition (*e.g.*, geography, diet, lifestyle) could have confounded the observed correlations between African ancestry and the relative abundances of *Enterococcaceae* and Otu00068. To split apart the effects of genetic ancestry and non-genetic factors, we fitted linear regression models using (arcsin square-root transformed) relative abundances of these microbes as dependent variables, and genetic PC1 and PC2, city of origin, sex, age, diet (calorie and fiber intakes) and lifestyle (physical activity levels, smoking status, and medicament consumption) as explanatory variables. We found that differences in microbial abundances were actually related to the city of origin (*Enterococcaceae*: F_4,426_ = 6.98, p<0.0001; Otu00068: F_4,426_ = 6.76, p<0.0001) and the smoking status (*Enterococcaceae*: F_1,426_ = 4.45, p = 0.04; Otu00068: F_4,426_ = 4.61, p = 0.03), being highest in Barranquilla, the city with the highest contribution of African ancestry, and nonsmokers (Figure S1). However, they did not relate to the host genetic ancestry after accounting for covariates (p>0.10 for genetic PC1 and PC2 for both taxa).

### Cardiometabolic health outcomes are better explained by gut microbiota composition than by host genetic ancestry

Considering that we found limited evidence of an association between gut microbiota and host genetic ancestry, we next examined whether gut microbes and the participants’ ancestral genetic composition each associated with variables related to cardiometabolic health, diet and lifestyle. The risk of disease was assessed through a summary measure—the cardiometabolic risk scale—which totaled *Z*-scores of waist circumference, triglycerides, fasting insulin, diastolic blood pressure and high-sensitive C reactive protein (hs-CRP). These variables informed about different conditions involved in cardiometabolic disease, namely central obesity, dyslipidemia, insulin resistance, hypertension and low-grade systemic inflammation, respectively.

Individuals with high values of the cardiometabolic risk scale were more likely to be male; to be of older age; to have low levels of high density lipoprotein (HDL) cholesterol; high levels of total cholesterol, low density lipoprotein (LDL) cholesterol, very low density lipoprotein (VLDL) cholesterol, and triglycerides; high levels of fasting glucose, glycated hemoglobin (HbA1c), fasting insulin and insulin resistance (HOMA-IR); high levels of hs-CRP, blood pressure and adiposity (BMI, waist circumference and body fat); and to regularly smoke and consume medications. In addition, these individuals were more likely to suffer of coronary heart disease, as assessed by the Framingham score [22]. While the cardiometabolic disease risk was not associated with genetic ancestry, diet intake or levels of physical activity, it was significantly associated with gut microbiota composition (Table 1).

**Table 1.**
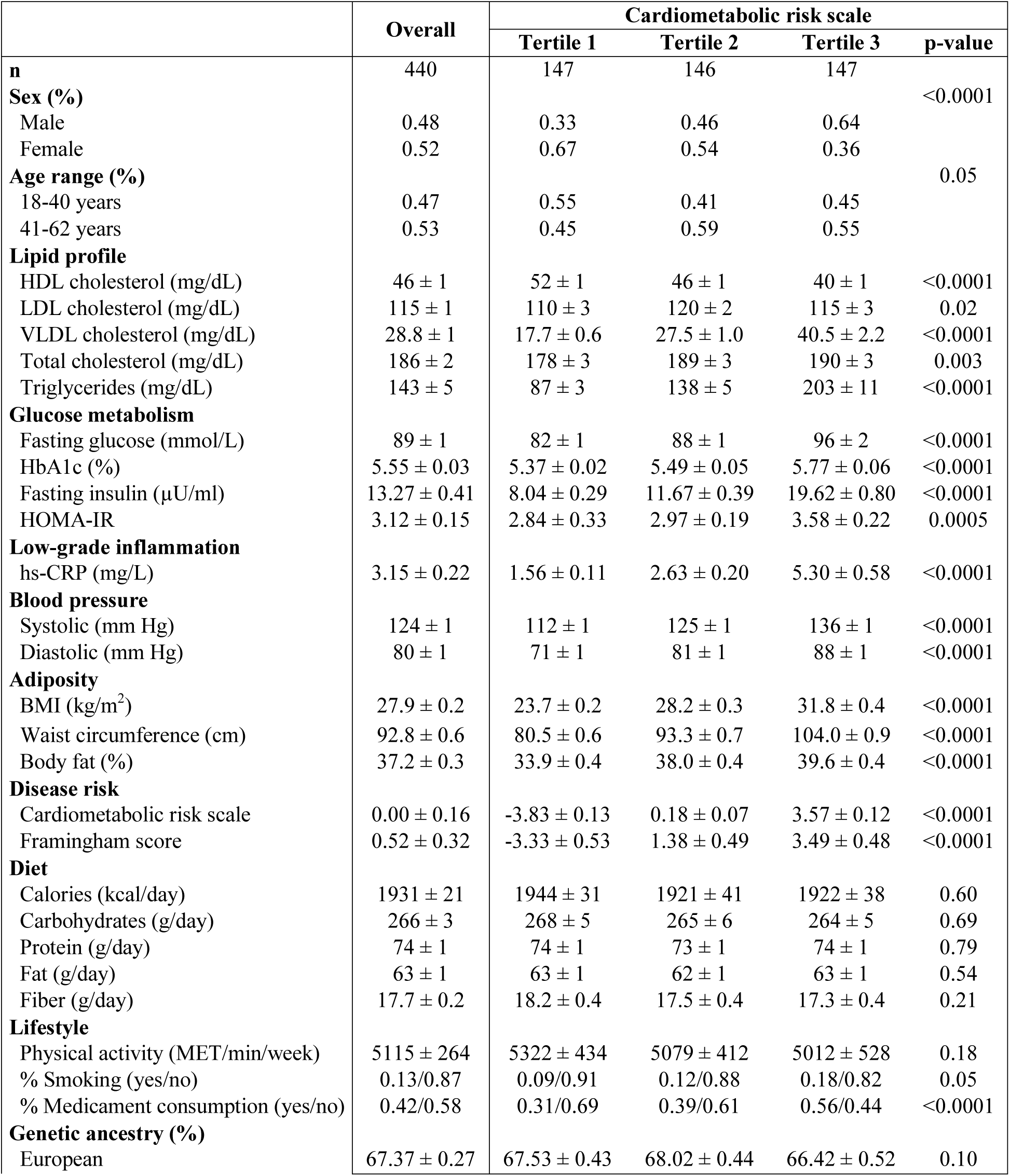
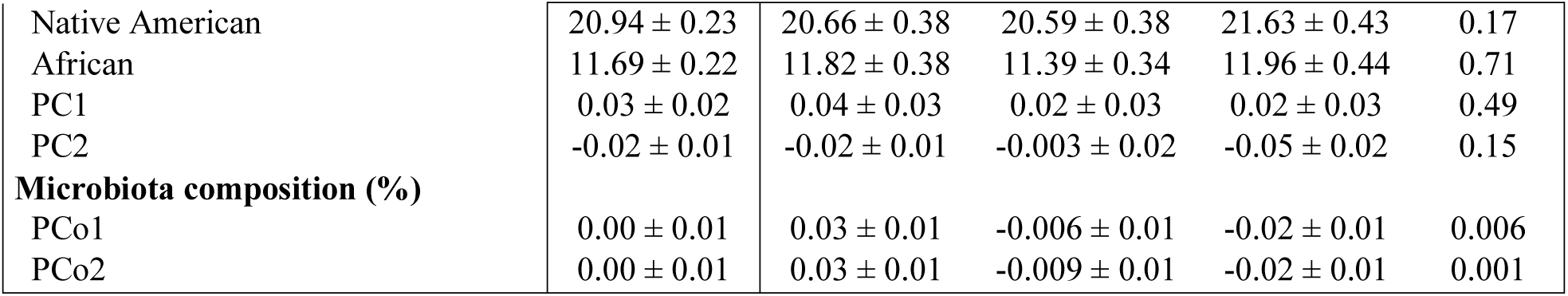
Characteristics of the study population. Variables presented overall and according to tertiles of the cardiometabolic risk scale. Data presented as mean ± SEM. *P*-values from ANOVA to the exception of sex, age range, smoking status and medicament consumption (chi-squared tests).

|  | Overall | Cardiometabolic risk scale |  |  |  |
| --- | --- | --- | --- | --- | --- |
|  |  | Tertile 1 | Tertile 2 | Tertile 3 | p-value |
| **n** | 440 | 147 | 146 | 147 |  |
| **Sex (%)** |  |  |  |  | <0.0001 |
| Male | 0.48 | 0.33 | 0.46 | 0.64 |  |
| Female | 0.52 | 0.67 | 0.54 | 0.36 |  |
| **Age range (%)** |  |  |  |  | 0.05 |
| 18-40 years | 0.47 | 0.55 | 0.41 | 0.45 |  |
| 41-62 years | 0.53 | 0.45 | 0.59 | 0.55 |  |
| **Lipid profile** |  |  |  |  |  |
| HDL cholesterol (mg/dL) | 46 ± 1 | 52 ± 1 | 46 ± 1 | 40 ± 1 | <0.0001 |
| LDL cholesterol (mg/dL) | 115 ± 1 | 110 ± 3 | 120 ± 2 | 115 ± 3 | 0.02 |
| VLDL cholesterol (mg/dL) | 28.8 ± 1 | 17.7 ± 0.6 | 27.5 ± 1.0 | 40.5 ± 2.2 | <0.0001 |
| Total cholesterol (mg/dL) | 186 ± 2 | 178 ± 3 | 189 ± 3 | 190 ± 3 | 0.003 |
| Triglycerides (mg/dL) | 143 ± 5 | 87 ± 3 | 138 ± 5 | 203 ± 11 | <0.0001 |
| **Glucose metabolism** |  |  |  |  |  |
| Fasting glucose (mmol/L) | 89 ± 1 | 82 ± 1 | 88 ± 1 | 96 ± 2 | <0.0001 |
| HbA1c (%) | 5.55 ± 0.03 | 5.37 ± 0.02 | 5.49 ± 0.05 | 5.77 ± 0.06 | <0.0001 |
| Fasting insulin (μU/ml) | 13.27 ± 0.41 | 8.04 ± 0.29 | 11.67 ± 0.39 | 19.62 ± 0.80 | <0.0001 |
| HOMA-IR | 3.12 ± 0.15 | 2.84 ± 0.33 | 2.97 ± 0.19 | 3.58 ± 0.22 | 0.0005 |
| **Low-grade inflammation** |  |  |  |  |  |
| hs-CRP (mg/L) | 3.15 ± 0.22 | 1.56 ± 0.11 | 2.63 ± 0.20 | 5.30 ± 0.58 | <0.0001 |
| **Blood pressure** |  |  |  |  |  |
| Systolic (mm Hg) | 124 ± 1 | 112 ± 1 | 125 ± 1 | 136 ± 1 | <0.0001 |
| Diastolic (mm Hg) | 80 ± 1 | 71 ± 1 | 81 ± 1 | 88 ± 1 | <0.0001 |
| **Adiposity** |  |  |  |  |  |
| BMI (kg/m<sup>2</sup>) | 27.9 ± 0.2 | 23.7 ± 0.2 | 28.2 ± 0.3 | 31.8 ± 0.4 | <0.0001 |
| Waist circumference (cm) | 92.8 ± 0.6 | 80.5 ± 0.6 | 93.3 ± 0.7 | 104.0 ± 0.9 | <0.0001 |
| Body fat (%) | 37.2 ± 0.3 | 33.9 ± 0.4 | 38.0 ± 0.4 | 39.6 ± 0.4 | <0.0001 |
| **Disease risk** |  |  |  |  |  |
| Cardiometabolic risk scale | 0.00 ± 0.16 | -3.83 ± 0.13 | 0.18 ± 0.07 | 3.57 ± 0.12 | <0.0001 |
| Framingham score | 0.52 ± 0.32 | -3.33 ± 0.53 | 1.38 ± 0.49 | 3.49 ± 0.48 | <0.0001 |
| **Diet** |  |  |  |  |  |
| Calories (kcal/day) | 1931 ± 21 | 1944 ± 31 | 1921 ± 41 | 1922 ± 38 | 0.60 |
| Carbohydrates (g/day) | 266 ± 3 | 268 ± 5 | 265 ± 6 | 264 ± 5 | 0.69 |
| Protein (g/day) | 74 ± 1 | 74 ± 1 | 73 ± 1 | 74 ± 1 | 0.79 |
| Fat (g/day) | 63 ± 1 | 63 ± 1 | 62 ± 1 | 63 ± 1 | 0.54 |
| Fiber (g/day) | 17.7 ± 0.2 | 18.2 ± 0.4 | 17.5 ± 0.4 | 17.3 ± 0.4 | 0.21 |
| **Lifestyle** |  |  |  |  |  |
| Physical activity (MET/min/week) | 5115 ± 264 | 5322 ± 434 | 5079 ± 412 | 5012 ± 528 | 0.18 |
| % Smoking (yes/no) | 0.13/0.87 | 0.09/0.91 | 0.12/0.88 | 0.18/0.82 | 0.05 |
| % Medicament consumption (yes/no) | 0.42/0.58 | 0.31/0.69 | 0.39/0.61 | 0.56/0.44 | <0.0001 |
| **Genetic ancestry (%)** |  |  |  |  |  |
| European | 67.37 ± 0.27 | 67.53 ± 0.43 | 68.02 ± 0.44 | 66.42 ± 0.52 | 0.10 |
| Native American | $20.94 \pm 0.23$ | $20.66 \pm 0.38$ | $20.59 \pm 0.38$ | $21.63 \pm 0.43$ | 0.17 |
| African | $11.69 \pm 0.22$ | $11.82 \pm 0.38$ | $11.39 \pm 0.34$ | $11.96 \pm 0.44$ | 0.71 |
| PC1 | $0.03 \pm 0.02$ | $0.04 \pm 0.03$ | $0.02 \pm 0.03$ | $0.02 \pm 0.03$ | 0.49 |
| PC2 | $-0.02 \pm 0.01$ | $-0.02 \pm 0.01$ | $-0.003 \pm 0.02$ | $-0.05 \pm 0.02$ | 0.15 |
| **Microbiota composition (%)** |  |  |  |  |  |
| PCo1 | $0.00 \pm 0.01$ | $0.03 \pm 0.01$ | $-0.006 \pm 0.01$ | $-0.02 \pm 0.01$ | 0.006 |
| PCo2 | $0.00 \pm 0.01$ | $0.03 \pm 0.01$ | $-0.009 \pm 0.01$ | $-0.02 \pm 0.01$ | 0.001 |

We verified these results by correlating variables summarizing genetic ancestry (PCA components) and gut microbiota (first two PCoA axes of weighted UniFrac) with biochemical profiles, blood pressure, adiposity, diet and physical activity. For this, we fitted linear models adjusted for the city of origin, sex, age, smoking status and medicament consumption, and calculated Spearman correlation coefficients between pairs of adjusted variables, so that correlations were independent of the aforementioned covariates. We found that the levels of blood insulin were negatively correlated with genetic PC2 (*i.e.*, individuals with higher non-European ancestries had higher insulin levels). On the other hand, microbiota PCoA axes were significantly associated with glucose metabolism (fasting glucose levels), hypertension (blood pressure), obesity (BMI and % body fat), central obesity (waist circumference), and fiber intake (Table 2).

**Table 2.** Multivariable-adjusted correlations between cardiometabolic outcomes, diet and physical activity with the host genetic ancestry and gut microbiota. Variables adjusted for the city of origin, age, sex, smoking status and medicament consumption. Spearman correlation coefficients (rho) and FDR-adjusted p-values (q-values) are shown. Values in bold highlight significant correlations.

|  | Genetic ancestry |  |  |  | Gut microbiota |  |  |  |
| --- | --- | --- | --- | --- | --- | --- | --- | --- |
|  | PC1 |  | PC2 |  | PCo1 |  | PCo2 |  |
|  | rho | q-value | rho | q-value | rho | q-value | rho | q-value |
| **Lipid profile** |  |  |  |  |  |  |  |  |
| HDL cholesterol | 0.03 | 0.82 | -0.02 | 0.91 | 0.06 | 0.67 | 0.02 | 0.91 |
| LDL cholesterol | 0.03 | 0.85 | -0.04 | 0.79 | 0.03 | 0.83 | -0.04 | 0.77 |
| VLDL cholesterol | -0.05 | 0.71 | -0.01 | 0.93 | -0.05 | 0.69 | -0.08 | 0.45 |
| Total cholesterol | 0.0001 | 0.99 | -0.05 | 0.69 | 0.04 | 0.79 | -0.05 | 0.69 |
| Triglycerides | -0.05 | 0.70 | -0.01 | 0.93 | -0.05 | 0.69 | -0.08 | 0.45 |
| **Glucose metabolism** |  |  |  |  |  |  |  |  |
| Fasting glucose | -0.05 | 0.70 | -0.01 | 0.92 | **-0.13** | **0.08** | -0.01 | 0.94 |
| HbA1c | 0.04 | 0.75 | -0.08 | 0.41 | 0.004 | 0.96 | 0.03 | 0.86 |
| Fasting insulin | -0.06 | 0.68 | **-0.13** | **0.05** | -0.11 | 0.21 | -0.07 | 0.54 |
| HOMA-IR | -0.02 | 0.91 | -0.04 | 0.74 | -0.10 | 0.27 | -0.03 | 0.83 |
| **Low-grade inflammation** |  |  |  |  |  |  |  |  |
| hs-CRP | -0.003 | 0.96 | -0.04 | 0.79 | -0.06 | 0.64 | -0.10 | 0.27 |
| **Blood pressure** |  |  |  |  |  |  |  |  |
| Systolic | -0.02 | 0.91 | 0.02 | 0.91 | **-0.13** | **0.05** | **-0.15** | **0.04** |
| Diastolic | 0.05 | 0.70 | 0.02 | 0.91 | -0.11 | 0.19 | **-0.16** | **0.03** |
| **Adiposity** |  |  |  |  |  |  |  |  |
| BMI | 0.01 | 0.93 | -0.10 | 0.27 | -0.09 | 0.31 | **-0.16** | **0.03** |
| Waist circumference | 0.005 | 0.96 | -0.05 | 0.69 | -0.08 | 0.45 | **-0.13** | **0.05** |
| Body fat | -0.03 | 0.79 | -0.06 | 0.64 | -0.06 | 0.67 | **-0.14** | **0.05** |
| **Diet** |  |  |  |  |  |  |  |  |
| Calories | 0.03 | 0.79 | 0.01 | 0.93 | -0.02 | 0.91 | -0.04 | 0.74 |
| Carbohydrates | 0.01 | 0.93 | 0.01 | 0.95 | 0.01 | 0.93 | -0.07 | 0.54 |
| Protein | 0.03 | 0.79 | 0.06 | 0.67 | -0.01 | 0.93 | -0.07 | 0.60 |
| Fat | 0.05 | 0.69 | -0.01 | 0.94 | -0.04 | 0.77 | 0.03 | 0.86 |
| Fiber | -0.04 | 0.74 | -0.03 | 0.79 | 0.01 | 0.93 | **-0.15** | **0.04** |
| **Lifestyle** |  |  |  |  |  |  |  |  |
| Physical activity | 0.02 | 0.88 | -0.08 | 0.45 | -0.02 | 0.91 | -0.01 | 0.95 |

We next examined the contributions of host genetic ancestry, gut microbiota and their interaction to explain variance in cardiometabolic disease risk using multivariable-adjusted linear models. Models were adjusted for the city of origin, sex, age, calorie and fiber intakes, levels of physical activity, smoking status and medicament consumption. Based on likelihood-ratio tests and the Akaike information criterion (AIC), we found that gut microbiota composition explained more variance in the risk of cardiometabolic disease than genetic ancestry (model including genetic ancestry: χ^2^ (2 df) = 4.41, p = 0.11, AIC = 2262; model including gut microbiota: χ^2^ (2 df) = 22.9, p<0.0001, AIC = 2243; model including genetic ancestry × gut microbiota interaction: χ^2^ (4 df) = 3.14, p = 0.53, AIC = 2248). Similar results were obtained for waist circumference, blood pressure and the Framingham coronary heart disease score (Table S4). The model that best explained variance in insulin levels was that considering genetic ancestry, whereas triglycerides levels were best explained by the ancestry × gut microbiota interaction. Neither genetic ancestry nor gut microbiota seemed to significantly contribute to explain variance in hs-CRP levels (Table S4).

Considering that gut microbiota associated with more health-related variables than genetic ancestry, we identified particular OTUs associated with cardiometabolic outcomes. For this, we analyzed the 100 most abundant OTUs and fitted quasi-Poisson generalized linear models (GLMs) on rarefied sequence counts, adjusting for the city of origin, sex, age, calorie and fiber intakes, physical activity, smoking status and medicament consumption. We calculated multivariable-adjusted Spearman correlation coefficients between OTU abundances and cardiometabolic outcomes, and obtained FDR-adjusted p-values. Independent of the aforementioned covariates, we found 19 OTUs significantly correlated with the cardiometabolic risk scale, eight with waist circumference, two with blood pressure and one with hs-CRP. No OTUs significantly correlated with triglyceride and fasting insulin levels or the Framingham coronary heart disease score. The relative abundances of nine OTUs, including *Escherichia coli, Atopobium, Gemmiger formicilis* and *Clostridiaceae SMB53*, among others, correlated with increased cardiometabolic disease risk, whereas 10 OTUs related to *Akkermansia muciniphila, Oscillospira, Methanobrevibacter* and *Christensenellaceae*, among others, correlated with lower disease risk (Figure 3).

**Figure 3.**
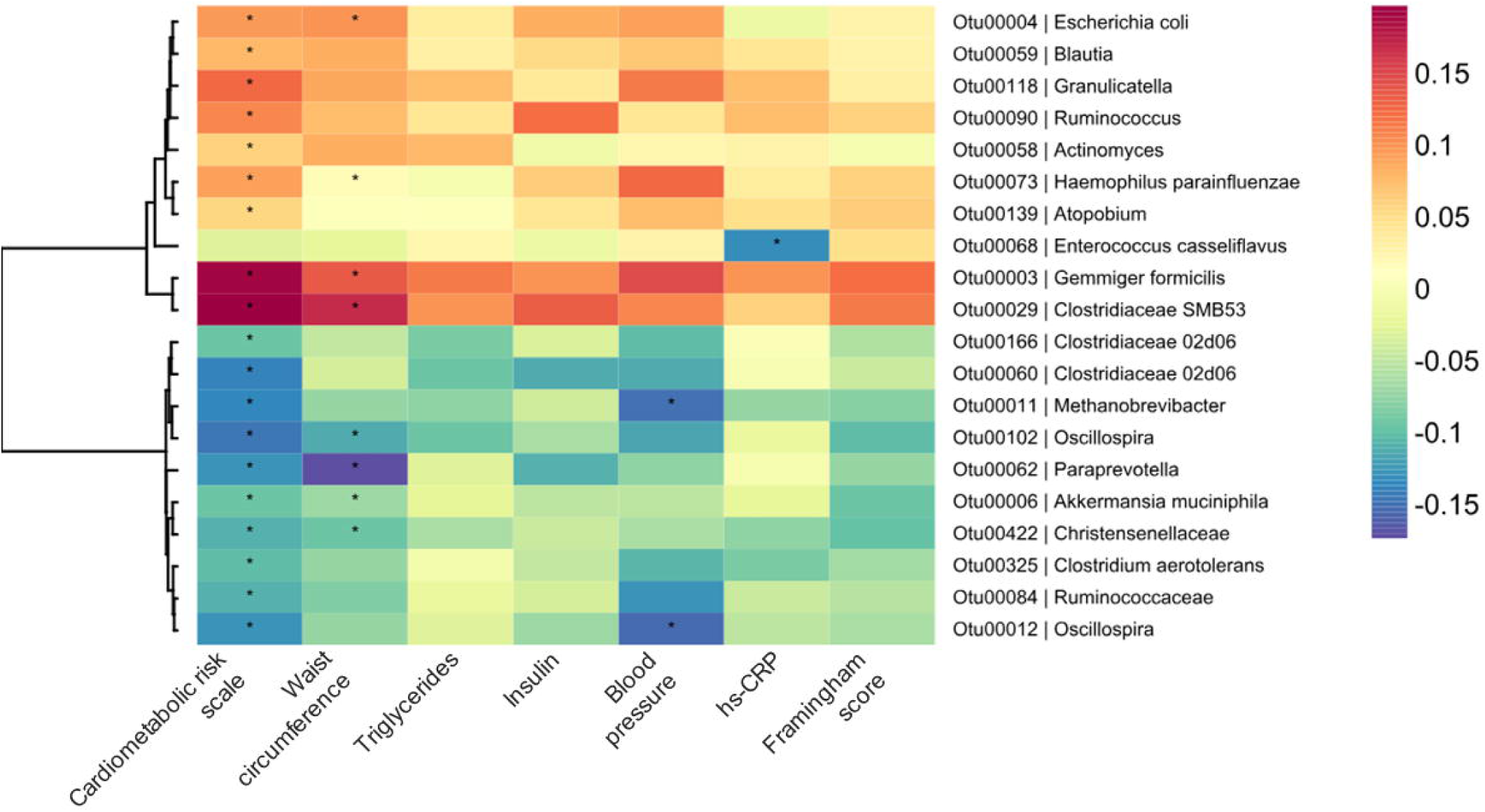
Heatmap showing the correlations between rarefied OTU abundances and multivariable-adjusted cardiometabolic outcomes. The dendrogram to the left was obtained by hierarchical Ward-linkage clustering based on correlation coefficients of the relative abundances of the 100 OTUs that had median abundances ≥0.01% across all participants. Correlations adjusted for the city of origin, sex, age, calorie and fiber intakes, physical activity, smoking status and medicament consumption. The color scale indicates Spearman correlation coefficients. FDR-adjusted p-values from quasi-Poisson generalized linear models are indicated (*=*q*<0.10).

## Discussion

Gut microbiota composition and the host genetic background have been each associated with human cardiometabolic health. However, the evidence associating the microbial community and the host genetic ancestry is sparse. We examined associations among host ancestry, gut microbiota and cardiometabolic health in a population with a history of recent, extensive admixture between Europeans, Native Americans and Africans [19]. Importantly, we quantified the levels of genetic admixture using ancestry informative markers (AIMs) located on most chromosomes, in opposition to self-reported ancestry [18,23]. While we found strong connections between gut microbiota and cardiometabolic health, the evidence associating these variables with the host genetic ancestry was limited.

The studied population had an admixed genetic composition typical of urban Latin American mestizos, with predominance of European, followed by Native American and African ancestries [19]. Overall, the contributions of each ancestral component followed a previously described geographic pattern, where inhabitants of the inner, Andean regions (Bogota, Medellin and Bucaramanga) had the highest European ancestry; those North and Northwest the lowest Amerindian ancestry (Medellin and Barranquilla); and those on the Caribbean and Pacific coasts (Barranquilla and Cali) the highest African ancestry [19,24]. These results confirmed that the city of origin was associated with the ancestral genetic composition of the studied population and that the panel of selected AIMs adequately replicated results from previous studies in Colombians.

Comparative cross-species studies indicate that hosts and symbionts have coevolved for millions of years [25,26], suggesting a heritable basis in this interaction. In agreement with this, recent studies in mice and humans suggest that gut microbiota is partly under the host genetic control [27–30], and particular microbes have been shown to be heritable [31]. However, the contribution of the host genetic ancestry to gut microbiota composition has been poorly studied, albeit it could be crucial to explain pervasive inter-population differences in this community [16,17].

We found a general lack of association between host genetic ancestry and gut microbiota composition in a mestizo population, agreeing with a recent study performed in Israel considering a variety of self-reported ancestries, including Ashkenazi, North African, Middle Eastern, Sephardi, Yemenite and admixed [18]. The only correlations detected in our study between genetic ancestry and gut microbiota came from *Enterococcaceae* and Otu00068 (*E. casseliflavus*), whose association with African ancestry was confounded by non-genetic factors, specifically the city of origin and cigarette consumption.

Multiethnic surveys have demonstrated that the origin of human populations contribute to the genetic predisposition to disease. Here, we found that individuals with higher Amerindian and African ancestries had higher blood insulin, independent of potential non-genetic confounders, including sex, age, the city of origin, diet and lifestyle. Studies in Mexican-Americans [32], US Native Americans [33] and Alaska Natives [34] have shown higher risk of type 2 diabetes in individuals of Amerindian ancestry. Likewise, Africans, African Americans and genetically-admixed individuals with high African ancestry have higher risk of this disease [35–38].

Our analyses, however, did not reveal further associations between the ancestral genetic composition and cardiometabolic health. This contrasts with recent studies performed in Colombians, which have shown associations between Native American ancestry and high triglyceride levels [39]; African ancestry, high blood pressure and high risk of type 2 diabetes [38,39]; European ancestry and low risk of type 2 diabetes [40]. One possible explanation for the general lack of association between genetic ancestry and health of our study could be the relative homogeneity in the individual proportions of European, Native American and African (Figure 1A). Unlike studies in which subjects with very dissimilar origin are compared, the individuals analyzed here were all mestizos with roughly similar admixed ancestry. The inclusion of individuals with more diverse genetic backgrounds could allow finding clearer health–ancestry associations [38,41,42].

An alternative, non-exclusive explanation could be that the correlations between genetic ancestry and health are actually small and require larger sample sizes and more AIMs to uncover them. For the former, we calculated the sample sizes needed to detect statistically significant differences in genetic ancestry among tertiles of the cardiometabolic risk scale, as reported in Table 1, and found that the statistical power of our study was indeed limited. For α = 0.05 and β = 0.80, we would have needed to enroll 860 individuals to detect statistically significant differences in European ancestry across levels of disease risk, 1108 for Native American, and 4849 for African. Concerning the number of evaluated AIMs, the power to detect statistical associations with a phenotype depends on the genome coverage of the evaluated genetic markers and on the basal linkage disequilibrium of the studied population. Although we only evaluated 40 AIMs, it was previously shown that as few as 30 AIMs allow getting an accurate representation of the ancestral genetic composition of Latin American populations [19]. In addition, our ancestry estimates at the population level were similar to those obtained in studies evaluating different set of variants [19] and population samples [39].

While the evidence associating genetic ancestry and health in our cohort was weak, the association between gut microbes and cardiometabolic outcomes was stronger. We found that the microbiota composition was a better explanatory variable of the risk of cardiometabolic disease than genetic ancestry, and informed about central obesity, hypertension and coronary heart disease. Further, we uncovered a list of particular OTUs associated with disease risk in the studied population. This included microbes that have been shown to be more abundant in patients with atherosclerotic disease, such as *E. coli* and *Atopobium* [11]; in type 2 diabetic patients, such as *Clostridiaceae SMB53* [43]; and in unhealthy obese individuals, such as *E. coli, Gemmiger formicilis* and *Clostridiaceae SMB53* [44]. On the other hand, microbes such as *A. muciniphila, Oscillospira, Methanobrevibacter* and *Christensenellaceae* have been shown to be associated to healthy cardiometabolic states [14,45–47].

Our study had several strengths, including a thorough sampling in several cities and an in-depth characterization of the studied cohort in terms of genetic ancestry, gut microbiota, cardiometabolic health outcomes and non-genetic factors associated with diet and lifestyle that allowed adjusting statistical models for potential confounding. However, for some comparisons we were limited by the sample size of our study, by the number of evaluated AIMs and by the fact that this was a cross-sectional study, so that we cannot distinguish cause and effect.

## Conclusions

Genetic-association studies have given great insights into the nature of complex phenotypes, such as cardiometabolic diseases. Accumulating evidence on non-genetic factors intimately linked to the host, such as the gut microbiota, has enriched this picture, demonstrating an intimate and complex connection between symbionts and human health. We here showed that the specific composition of gut microbes has a dramatic effect on disease risk, ultimately opening a promising avenue to ameliorate human health through targeted modulation of the microbial community. Our results, and those recently obtained in a different population [18], suggest that modulation strategies could be applied regardless of the genetic ancestry of the intervened population.

## Methods

### Study population

We enrolled 441 mestizo adult men and women, living in the cities of Bogota, Medellin, Cali, Barranquilla and Bucaramanga (Colombia, South America) between July and November 2014. The national census indicates that these cities contribute about 30% of the Colombian population. Participants were enrolled in similar proportions according to the city of residence (19% Bogota, 22% Medellin, 20% Cali, 20% Barranquilla and 18% Bucaramanga), BMI (31% lean, 39% overweight and 30% obese), sex (48% male, 52% female), and age range (47% 18-40 years, and 53% 41-62 years). We excluded underweight participants (*i.e.*, BMI<18.5 kg/m^2^), pregnant women, individuals who had consumed antibiotics or antiparasitics in the three months prior to enrollment, and individuals diagnosed with neurodegenerative diseases, current or recent cancer (<1 year), and gastrointestinal diseases (Crohn’s disease, ulcerative colitis, short bowel syndrome, diverticulosis or celiac disease).

### Genotyping of ancestry informative markers (AIMs)

The ancestral genetic composition of participants was assessed through a panel of 40 AIMs located on most chromosomes, chosen for having strong differences in allele frequency between European, Native American and African populations, and to be unlinked (Table S1). The selected AIMs have been previously used [24,39,48,49]. Of these, 34 corresponded to insertion/deletion variants (INDELs) and six to single nucleotide polymorphisms (SNPs). Primers and PCR conditions followed specific protocols for each AIM. For INDELs, genotypes were resolved with 1.5-2.0% agarose gel electrophoresis if the variant was >10 bp, otherwise with capillary electrophoresis in an ABI PRISM 3100 Genetic Analyzer (Applied Biosystems, Foster City, CA). SNPs were genotyped with PCR-RFLP and resolved with 2.5-3.0% agarose gel electrophoresis.

### Analysis of host genetic ancestry

The host genetic ancestry was analyzed as follows: genotypes for each AIM served to calculate the observed and expected allelic and genotypic frequencies, to test the Hardy-Weinberg equilibrium with an exact test [50], and to estimate overall population structure (*F*_st_) using the Weir and Cockerham estimator [51]. The standard error and 95% confidence intervals of this estimator were calculated by jackknifing and bootstrapping over loci, respectively. Population-genetic analyses were performed with GenePop [52] and FSTAT 2.9.3 [53]. Afterwards, we tested isolation by distance by correlating genetic (*F*_st_/(1-*F*_st_)) and (log-transformed) geographic distance matrices using a Mantel test, as implemented in the ecodist package of R [54], with 10,000 permutations and 10,000 bootstrap iterations for calculating confidence intervals.

Next, a hidden Markov model approach was used to infer the individual genetic contributions of European, Native American and African ancestries using ADMIXMAP 3.7. [55]. This method models individual admixture using genotypic information for all individuals and AIMs, the AIM’s physical position in the chromosome and the frequency of the largest allele in parental populations. Allelic frequencies in the parental populations were previously reported for Europeans (Spain, Germany, England, Ireland), Native Americans (Maya, Pima and Puebla) and Africans (Nigeria, Sierra Leone, Central African Republic, African-American and Afro-Caribbean) [56,57]. The parameters used for running ADMIXMAP were: 40 loci, 440 diploid individuals, 250,000 iterations with a burn-in of 10,000 iterations, and a model of three populations.

The proportions of European, Native American and African ancestries were compared across the five cities from which our participants originated, BMI (lean, overweight, obese), sex (male, female) and age range (18-40, 41-62 years) with ANOVA, after verifying homoscedasticity with the Fligner-Killeen test. Where necessary, data were transformed with natural logarithm for unbounded variables, or arcsin square root for proportions. We also performed robust principal components analysis (PCA) for compositional data with the individual proportions of the three genetic ancestries using the robCompositions package of R [58]. For this, the compositional dataset was transformed using the isometric log ratio, and a PCA was afterwards performed. PC1 and PC2 components were compared across cities, BMI, sex and age range using ANOVA.

### Gut-microbiota characterization

Detailed laboratory and bioinformatic procedures can be found elsewhere [59]. Briefly, each participant collected a fecal sample from which the total microbial DNA was extracted using the QIAamp DNA Stool Mini Kit (Qiagen; Hilden, Germany). The V4 region of the 16S rRNA gene was amplified with primers F515 and R806, sequenced with Illumina MiSeq v2, and processed as previously described [59].

The gut microbiota was analyzed at the whole community level using principal coordinate analysis (PCoA) based on weighted UniFrac distances. These distances were computed on rarefied sequence counts (3667 reads/sample) with the GUniFrac package of R [60], and compared across cities, BMI, sex and age range with permutational multivariate analysis of variance using distance matrices (PERMANOVA), as implemented in the Vegan package of R [61]. Microbiota analyses were also performed at the phylum, class, order, family, genus and species level, as well as at the OTU level. For taxonomy-based tests, we calculated the relative abundance of microbial sequences classified at all taxonomic ranks according to the Greengenes 13_8_99 taxonomy [62]. At the OTU level, we grouped sequences at 97% identity using the average neighbor algorithm [63], and extracted the OTUs that had median relative abundances ≥0.01% across all samples. The latter procedure guaranteed that the majority of sequences was analyzed (∼80% of total reads) and minimized the impact of sequencing artifacts.

### Cardiometabolic health, diet and lifestyle

We measured several variables that might interact with both gut microbiota and the host genetic ancestry. These included health-related variables (blood chemistry, blood pressure and adiposity), diet intake (calories, macronutrients and dietary fiber) and lifestyle (physical activity, smoking status, medicament consumption). Detailed information about measurement of these variables is presented elsewhere [44]. Briefly, blood biochemical variables, including HDL, LDL, VLDL, total cholesterol, triglycerides, fasting glucose, HbA1c, fasting insulin, and hs-CRP, were measured using standard techniques routinely used in a clinical laboratory (Dinámica IPS, Medellin, Colombia). Blood insulin served to calculate the insulin resistance index using the homeostasis model assessment (HOMA-IR). The systolic and diastolic blood pressures were measured in mm Hg with a Rossmax AF701f digital tensiometer (Berneck, Switzerland). Adiposity was assessed through BMI (weight (kg)/height squared (m^2^)), waist circumference (cm) and percentage body fat (calculated with the thicknesses of four skinfolds: biceps, triceps, subscapular and ileocrestal).

To assess the risk of cardiometabolic disease, we constructed a summary scale—the cardiometabolic risk scale—by summing *Z*-scores of waist circumference, triglycerides, fasting insulin, diastolic blood pressure and hs-CRP (*Z* = [*x*-*µ*]/*δ*, where *µ* is the population mean and *δ* is the standard deviation of the population). Variables were log-transformed to adjust to a normal distribution before obtaining *Z*-scores. These variables were chosen because they informed about different conditions involved in cardiometabolic disease: central obesity, dyslipidemia, insulin resistance, hypertension and low-grade systemic inflammation, respectively. In addition, we calculated the Framingham coronary heart disease score [22] using sex, age, diabetes status, smoking status, blood pressure, HDL and total cholesterol as predictor variables. Since the Framingham score did not consider individuals younger than 30 years, these were given the lowest age score (−1).

Daily intakes of macronutrients (g/day of carbohydrates, protein and fat), dietary fiber (g/day) and calories (kcal/day) were estimated with 24-hour dietary recall interviews [64]. Physical activity (number of metabolic equivalents per minute per week: MET/min/week) with the short form of the International Physical Activity Questionnaire [65]. Smoking and medicament consumption were self-reported in specific questionnaires. For the latter, we considered all drugs taken by participants on a regular basis during the three months prior to enrollment, to the exception of over-the-counter vitamin and mineral supplements, phytotherapeutics and contraceptives. All measurements and questionnaires were performed by trained personnel.

### Associations of host genetic ancestry, gut microbiota and cardiometabolic health

The direct association between host genetic ancestry and microbiota composition was assessed with Procrustes analyses. These were performed to examine, on one hand, the correlation between the weighted UniFrac distance matrix and the matrix of individual proportions of European, Native American and African; and, on the other hand, the correlation between the first two PCoA axes of the microbiota analysis and the PCA components of genetic ancestry. In both cases, microbiota matrices were set as targets and genetic ancestry matrices as those to be rotated and scaled. Statistical significance was determined using 10,000 permutations.

We also explored associations between genetic ancestry and microbiota composition at the phylum, class, order, family, genus, species and OTU levels. In these cases, we correlated the relative abundance of each microbial group with the individual proportions of European, Native American and African, as well as with the two genetic PCA components, using Spearman correlation tests; p-values were adjusted for multiple comparisons (FDR) using the Benjamini-Hochberg method.

To dissect the effects of genetic ancestry and non-genetic factors on the abundance of particular groups of microbes, we fitted linear regression models in which the relative abundance of each microbial group was modeled in function of genetic ancestry (PCA components), city of origin, sex, age, diet (calorie and fiber intakes) and lifestyle (physical activity levels, smoking status, and medicament consumption). In these cases, relative abundances were arcsin square-root transformed.

We next investigated associations of the host genetic ancestry and gut microbiota composition with cardiometabolic health. For this, we divided the cardiometabolic risk scale by tertiles (low, middle and high risk) and tested differences among them for each variable using ANOVA and chi-square tests. Where necessary, variables were appropriately transformed as mentioned above.

Afterwards, we correlated variables informing about genetic ancestry (PCA components) and gut microbiota (first two PCoA axes of weighted UniFrac) with cardiometabolic outcomes, diet and physical activity. For this, we fitted linear models adjusted for the city of origin, sex, age, smoking status and medicament consumption, calculated Spearman correlation coefficients and obtained FDR-adjusted p-values for all pairs of adjusted variables.

To examine the contributions of host genetic ancestry, gut microbiota and their interaction in explaining cardiometabolic disease risk, we fitted several linear models. The basic model included the city of origin, sex, age, calorie and fiber intakes, levels of physical activity, smoking status and medicament consumption. We then evaluated alternative models including genetic ancestry (PCA components), gut microbiota (first two PCoA axes of weighted UniFrac) and the genetic ancestry × gut microbiota interaction. The first two alternative models were each compared against the basic model, the latter model was compared against the best preceding model. We obtained log-likelihoods of all models and evaluated their changes with likelihood ratio tests. Model selection was based on AIC. Models were fitted for the cardiometabolic risk scale, for individual variables adding up to this scale and for the Framingham coronary heart disease score.

Finally, we identified particular OTUs associated with cardiometabolic outcomes by fitting quasi-Poisson GLMs on rarefied sequence counts, adjusting for the city of origin, sex, age, calorie and fiber intakes, physical activity, smoking status and medicament consumption. The residuals of these GLMs were then correlated with cardiometabolic outcomes using Spearman correlation coefficients and FDR-adjusted p-values.

### Power calculation

We performed statistical power calculations to determine the sample sizes required to observe significant differences in genetic ancestry among tertiles of cardiometabolic disease risk, using the pwr package of R [66]. For this, we set the significance level (α = 0.05) and statistical power (β = 0.80), calculated the within-group variance for each ancestral genetic composition (proportions of European, Native American and African), and effect sizes (*f*). The latter were calculated using:

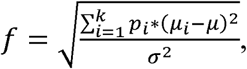

where *k* is the number of groups, *p*_*i*_ = *n*_*i*_/*N, n*_*i*_ is the number of observations in group *i, N* is the total number of observations, *µ*_*i*_ is the mean in group *i, µ* is the grand mean, and *σ*^2^ is the error variance within groups [67].

## List of abbreviations

AIC: Akaike information criterion
AIM: ancestry informative marker
BMI: body mass index
FDR: false discovery rate
GLM: generalized linear model
GWAS: genome-wide association study
HbA1c: glycated hemoglobin
HDL: high density lipoprotein cholesterol
HOMA-IR: homeostasis model assessment-insulin resistance
hs-CRP: high-sensitive C reactive protein
INDEL: insertion/deletion
LDL: low density lipoprotein cholesterol
MET/min/week: number of metabolic equivalents per minute per week
OTU: operational taxonomic unit
PCA: principal component analysis
PCoA: principal coordinate analysis
PCR-RFLP: polymerase chain reaction-random fragment length polymorphism
PERMANOVA: permutational multivariate analysis of variance
SNP: single nucleotide polymorphism
VLDL: very low density lipoprotein cholesterol.

## Declarations

### Ethics approval and consent to participate

The study followed the principles of the Declaration of Helsinki and had minimal risk according to the Colombian Ministry of Health (Resolution 8430 of 1993). Written informed consent was obtained from all the participants prior to the beginning of the study. The study was approved by the Bioethics Committee of SIU—University of Antioquia (act 14-24-588 dated May 28, 2014).

### Consent for publication

Not applicable.

### Availability of data and material

Raw 16S rRNA gene reads were deposited at the short read archive (BioProject PRJNA417579). Participants’ genetic and phenotypic data were deposited at the database of Genotypes and Phenotypes (dbGaP) (accession XXXX). The employed R code used in this paper is available at Github (https://github.com/jsescobar/microbiota_ancestry_health).

### Authors’ contributions

JSE and GB conceived the study. SJG-C, ELO-V, WR and GB obtained human genotypes and performed host genetic analysis. JdlC-Z, EPV-M and JSE obtained microbial DNA and performed gut microbiota analysis. SJG-C and JSE drafted the manuscript with contributions of all authors.

## Acknowledgments

We thank the participants who took part in the study. We are indebted to Roberto A. Jiménez and Luisa F. Mesa for their contributions during this project, and to GENMOL and Vidarium staff for their contributions during field and laboratory work. We are grateful to EPS SURA and Dinámica IPS for their support throughout the study. Some authors of this work collaborate through the Microbiome & Health Network.

